# Assessing pollinator community recovery in restored agroecosystems using the recovery debt framework

**DOI:** 10.64898/2026.05.08.723832

**Authors:** Domingo Cano, Antonio J. Pérez, Carlos Martínez-Núñez, Rubén Tarifa, Teresa Salido, Carlos Ruiz, José E. Gutiérrez, Julio M. Alcántara, Pedro J. Rey

**Affiliations:** Departamento de Biología Animal, Biología Vegetal y Ecología. Universidad de Jaén, Jaén, Spain; Estación Biológica de Doñana (EBD-CSIC), Sevilla, Spain; Estación Experimental de Zonas Áridas (EEZA-CSIC), Almería, Spain; SEO/BirdLife. Oficina del LIFE Olivares Vivos, GEOLIT, Mengíbar, Spain; Instituto Interuniversitario de Investigación del Sistema Tierra de Andalucía. Sede de la Universidad de Jaén, Jaén, Spain

**Keywords:** Agro-environmental measures, Ecological restoration, Ecological contrast, Ecosystem complexity, Olive groves, Permanent crops, Wild bees, Widlife-friendly farming

## Abstract

Recovery debt (RD) quantifies the interim deficit of biodiversity and function during the recovery process after disturbance. Unlike typical recovery indices derived from data on experimental-control comparisons, RD further considers the target (reference) biodiversity level, modelling the rate at which it is approached over time. However, the application of the RD approach to active restoration has not been explicitly implemented to date.

Here, we extend the RD framework to evaluate active ecological restoration in agricultural systems, defining the onset of recovery as the shift from intensive to wildlife-friendly management. We applied this approach to assess short-term pollinator recovery in 14 olive groves across a gradient of farming intensification and landscape complexity in southern Spain. Restoration actions included adopting low-intensity ground cover management and actively restoring field margins. At one, three, and five years post-restoration, we assessed community responses by quantifying bee abundance, species richness, plant–bee network properties, and flower visitation rates. Reference systems were defined by olive groves in complex landscapes with low-intensity herb cover management and organic farming practices.

Following restoration, the RD of bee abundance decreased from 71% to 55%. We found no significant effects of pre-intervention agricultural management on RD. Instead, across sites, the reduction of the RD (i.e., recovery) of bee abundance, richness, network connectance and flower visitation rate was strongly mediated by the availability of high-quality semi-natural areas in the surrounding landscape and by the ecological contrast created by restoration interventions at both the farm and floral patch levels. RD for other network metrics showed no significant pattern of variation.

Our study demonstrates that wildlife-friendly management and targeted habitat restoration can rapidly reduce recovery debt for bee abundance and function in permanent agroecosystems. However, the recovery of more complex interaction-network properties likely requires longer timescales.

## 1- Introduction

Landscape simplification by land conversion to agriculture and intensified farming practices are significant contributors to the global decline of pollinators (Potts et al., 2010; Raven & Wagner, 2021; Tilman et al., 2001). These factors directly affect the availability of nesting and floral resources, thus depauperating pollinator communities (Habel et al., 2019; Kennedy et al., 2013). Given the crucial role of pollinators in maintaining terrestrial ecosystems through the pollination function (Hagen Melanie & Kraemer Manfred, 2010; Klein et al., 2007), it is imperative to integrate agricultural production with pollinator conservation efforts. Ecological restoration emerges as a powerful approach to maintaining ecosystem services and biodiversity in agricultural landscapes (Rey Benayas & Bullock, 2012; Wade et al., 2008), and it is an essential idea emerging from the 2022 Kunming-Montreal Convention of Biological Diversity and the European Nature Restoration Law, among others. Indeed, the benefits of ecological restoration and wildlife-friendly practices for pollinators and the pollination service in agricultural areas are well described in the scientific literature (e.g., Bengtsson et al., 2005; Gaspar et al., 2022; Kovács-Hostyánszki et al., 2017; M’Gonigle et al. 2015).

For any specific group of organisms and/or community, the recovery process is not immediate, and its success depends on many factors (Moreno-Mateos et al., 2015). Similar to the concept of extinction debt, where species loss is delayed after environmental disturbance (Figueiredo et al., 2019), there is also a delay in biodiversity and ecosystem function recovery after restoration, or after the disturbance has ceased (Moreno-Mateos et al., 2017). This delay is called recovery debt (RD), defined as the interim (or temporary) deficit of biodiversity and functions occurring until the recovery process is completed (Moreno-Mateos et al., 2017).

Unlike typical recovery indices from data on experimental versus control plots (for example, before-after control impact designs, see Lengyel et al., 2023; see also Kremen & M’Gonigle 2015 and M’Gonigle et al., 2015, for the specific case of wild bees), RD further considers how much biodiversity we can recover (the reference value), and models how far we are from such value at each stage of the recovery process, the rate at which this reference value is approached and the time needed to reach it. This may be done through repeated measurements of biodiversity through time after disturbance has ceased or the restoration actions are set in place (Fig. 1).

**Figure 1.**
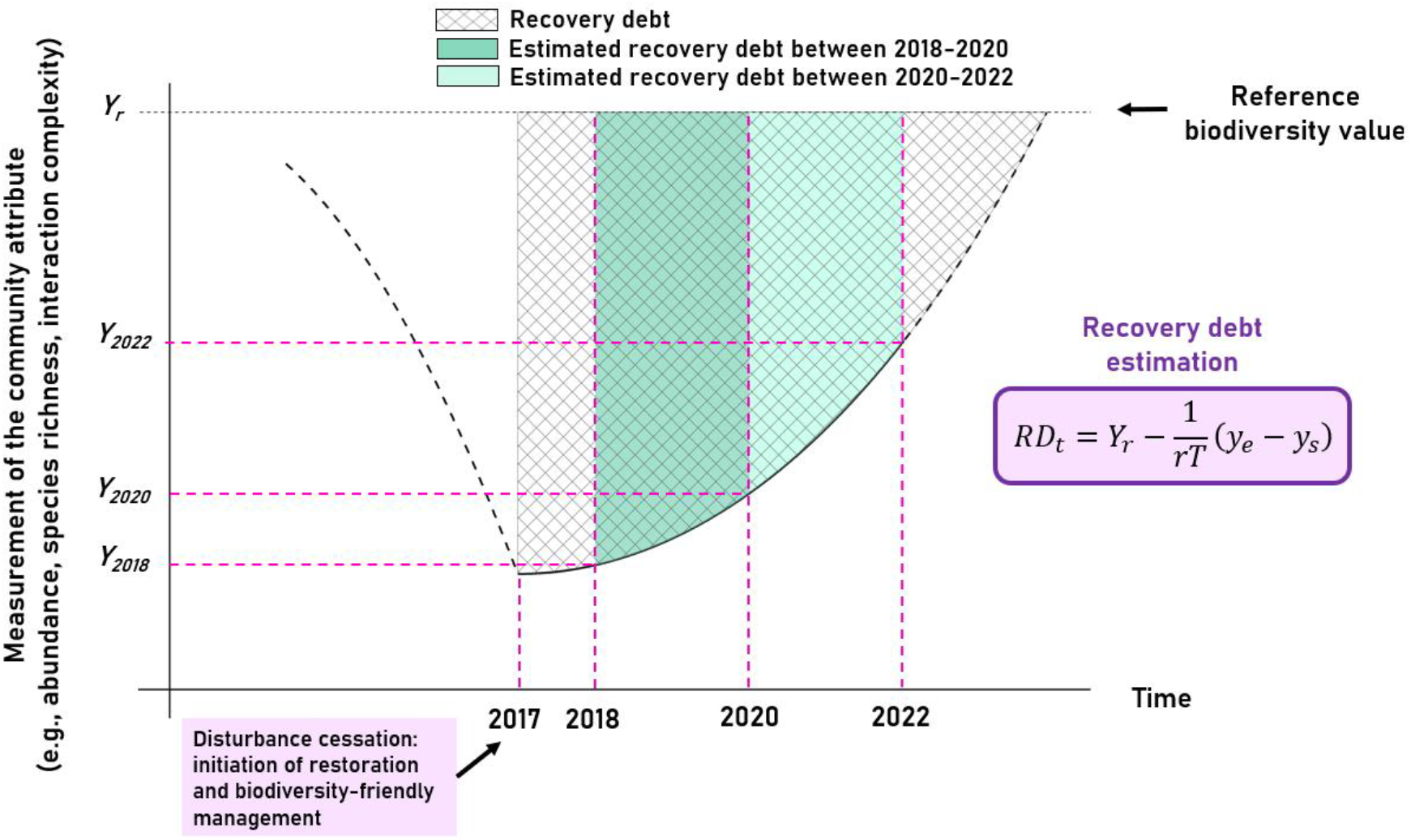
Measurement of the recovery debt (RD) in an agroecosystem, adapted from Moreno-Mateos et al. (2017). The figure illustrates the recovery process of a community attribute (e.g., species abundance or richness, ecosystem function, or interaction network descriptor) after the implementation of restoration actions and the approach through time to a reference value (i.e., the value of a well-preserved reference community representative of the same agroecosystem). Because the recovery is a dynamic process we can better estimate it through modelling the change in the community attribute by a repeated measure through time after restoration interventions, which will allow estimating the rate of recovery (rate at which the reference value is approached), the time needed for recovery and the amount of the attribute that still remains to be recovered at a given time or within a time interval. RD represents the interim deficit of the attribute during recovery, that is, the remaining distance to the reference value (*Y*_*r*_ – *Y*_*t*_) at a given time, or the area between the recovery curve and the reference level over a time interval. Following Moreno-Mateos (2017), this area may be calculated through the equation indicated in the figure, assuming an exponential rate of recovery and setting it as a per-annual rate (RD_t_). Thus, this figure illustrates the idea of RD and how it is calculated. The area with the grid pattern represents the change over time in the extent to which the ecosystem attribute remains reduced after disturbance cessation (in this case, the cessation of intensive farming and the initiation of restoration and biodiversity-friendly management in 2017). Dashed lines on the left and right sides of the curve represent, respectively, the presumed deterioration of the community attribute by agricultural disturbance and the expected exponential progression of recovery beyond our measures until it reaches the reference value. In this study, community attributes were measured in 2018, 2020, and 2022, allowing estimation of RD_t_ for the intervals 2018–2020, 2020–2022, and 2018–2022, though only species abundance and richness were measured at all three time points.

In the extinction debt approach, the rate and delay with which species loss occurs in response to disturbance depends on the community functional response (i.e., life-history traits) (Kuussaari et al., 2009), as well as on the intensity and frequency of the disturbance (Claudino et al., 2015). Similarly, recovery after restoration may also depend on these factors. Pollinating insects have short generation lengths, high mobility, and great ability to track resources in changing environments, and therefore they might respond more quickly than other groups to restoration actions (Brown et al., 2024; Kuussaari et al., 2009; Sexton & Emery, 2020). As a consequence, a relatively quick shortening of the RD linked to their process of recovery is expected. However, ecosystem recovery rates vary depending on the complexity of the ecosystem component assessed, with slower recovery rates as complexity increases (Moreno-Mateos et al., 2020). For example, theoretically, the recovery of species interactions after restoration interventions is slower than the recovery of species richness or abundance, as it requires more specific conditions such as encounter probability, the presence of species with matching traits, and phenological synchrony (Moreno-Mateos et al., 2020; Valiente-Banuet et al., 2015). Despite this, empirical studies have shown that plant-pollinator interactions also respond rapidly to restoration actions due to the adaptability of certain pollinators acting as generalists (Forup et al., 2008; Noreika et al., 2019; Traveset et al., 2024). Indeed, highly generalist pollinators have been observed to colonise restored sites at high rates (Ponisio et al., 2019). Therefore, to gain a more comprehensive knowledge about the degree of ecosystem degradation and recovery, we need to estimate RD over time through a series of attributes representing a gradient of ecosystem complexity (i.e., abundance and richness of species, ecosystem functions and ecological networks). Similarly, estimating RD after active restoration throughout gradients of disturbance (for example, gradients of agricultural intensification) would help us to assess the extent to which the disturbances linked to agricultural management affect the success of certain practices in promoting the recovery of biodiversity over time, an issue virtually unexplored.

Studies assessing the magnitude of the RD are still scarce, with the majority being meta-analyses primarily focused on spontaneous or quasi-spontaneous recovery across types of disturbances and ecosystems (e.g., Hillebrand & Kunze, 2020; Mori et al., 2020; Senf & Seidl, 2022). Here, we extend the RD concept to active ecological restoration in agroecosystems, considering disturbance cessation as the time of shifting from more intensive to wildlife-friendly practices (Fig. 1). Specifically, we applied this approach to assess short-term (5 years) recovery of pollinator biodiversity and function in 14 olive groves of Andalusia, southern Spain, across a gradient of landscape complexity. This was achieved by transitioning from their original farming practices to less intensive farming through decreases in the intensity of the herb cover management and/or use of pesticides, along with the implementation of active restoration measures. Olive groves, being the most widespread permanent woody crop in Europe and mainly growing within one of the Earth’s biodiversity hotspots (Martínez-Núñez et al., 2025), play a key role in pollinator conservation despite not being an insect-pollinated crop (Cano et al., 2022; Dellapiana et al., 2025; Martínez-Núñez et al., 2020a; Potts et al., 2006; Rey et al., 2019; Tscheulin et al., 2011). We assess their RD following active restoration actions and determine how RD is affected, in the short term, by the original farming practices of the olive groves, the magnitude of ecological change caused by restoration interventions (i.e., ecological contrast of the AES, *sensu* Marja et al., 2019, as a moderator of biodiversity recovery) and the landscape heterogeneity. This assessment is based on a set of target variables representing different levels of complexity in the pollinator community: abundance and richness of wild bees (hereafter “bees”), pollination function (using flower visitation rates as a proxy), and plant-bee interaction network metrics.

More specifically, we aim to i) evaluate how the RD is linked to the recovery of different properties of the bee communities over time since the start of restoration interventions in the olive farms; and ii) determine how RD depends on the original farming intensity, the heterogeneity at landscape and floral patch scales, and the magnitude of change (ecological contrast) driven by restoration interventions. We specifically expect the RD to decrease quicker for bee abundance and flower visitation rates than for richness or interaction network properties. Additionally, at early stages of recovery (the five-year post-intervention evaluated here), we anticipate a lower RD in olive groves that had environmentally friendly pre-intervention farming management (i.e., low-intensity herb cover management and organic farming). We further expect a lower RD as a function of the increased availability of semi-natural areas suitable for bees in the landscape and/or the ecological contrast caused by interventions aimed at benefiting pollinators.

## 2- Materials and Methods

### Study system

This study was conducted in the olive grove landscapes of Andalusia (southern Spain), the region with the largest cultivation area of this crop globally (> 1.5 million hectares). We selected 14 olive farms, which span 293 km apart, within the geographical extremes located at 5°28′18″W, 36°59′31″N and 2°38′41″W, 38°23′49’’N (Appendix S1: Fig. S1). These farms are part of the restoration program of the LIFE project “Olivares Vivos” (LIFE20 AT/ES/001487, European Commission) to demonstrate the effectiveness of biodiversity-friendly practices and nature restoration interventions to recover biodiversity. Before the interventions, these olive farms had different intensities of herb cover management (high-intensity vs. low-intensity) and use of pesticides (non-organic vs. organic) and were distributed across a gradient of landscape complexity. High-intensity management involves permanently removing the ground herb cover with the application of pre- and post-emergence herbicides or through deep and recurrent ploughing (in some cases, both coupled). Conversely, low-intensity management consists of maintaining the herb cover during most of the year, removing it only in late spring by mechanical mowing or livestock grazing (typically sheep). Regarding the use of pesticides, non-organic farms employ synthetic agrochemicals (insecticides, herbicides, and fertilisers), whereas organic farming does not allow their use. See Appendix S1: Table S1 for detailed information about the landscape and management characteristics of the olive groves of this study.

In the last third of 2016 and during 2017, all the study farms were subjected to diverse restoration actions aimed at recovering biodiversity, like the revegetation of boundaries and field margins with woody plants, the establishment of herbaceous patches by seed sowing, and the installation of artificial ponds and nests with cavities for solitary bees. Appendix S1 (Table S2) provides detailed information about the magnitude of change of each of the recovery measures implemented. Moreover, at the farm level, those olive farms that initially had high-intensity herb cover management switched to low-intensity practices, aimed at increasing the diversity and cover of herbaceous plants. Therefore, to estimate the RD of bees and their function (see below), we consider 2017 as the year of cessation of disturbance on all farms.

In this study system, we monitored bee abundance, diversity, and interactions with flowers at three different times after interventions: 2018, 2020, and 2022 (one, three, and five years post-restoration interventions, respectively) to estimate the RD. In each olive farm, we monitored two 10 m^2^ floral herb patches. We further quantified farm-level herbaceous cover and diversity before and after interventions, the surrounding landscape complexity, and the floral characteristics of the pollinator sampling patches (see details on predictor estimation in the following sections).

This allows considering different types of predictors of the RD: (i) predictors of the original management intensity of the farm (see methodological details below); (ii) predictors of the pre-intervention heterogeneity at landscape and floral patch scales; and (iii) predictors of the magnitude of change mainly concerning enhancement in floral cover and richness, measured at both farm and patch scales.

### Response variables: wild bee abundance, diversity, and interaction with flowers

We calculated the RD associated with the recovery of the bee community structure and function in olive groves post-restoration. We employed three sets of target variables: abundance and richness, as basic descriptors of community structure, and plant-bee interaction variables derived from interaction networks, as complex indicators of community assemblage and quality of pollination function.

Data collection consisted of surveys in twenty-eight 10 m^2^ multi-floral patches within the olive crop matrix (two patches per olive farm). We carried out two sampling rounds per olive farm between April and June each year, with approximately one month between rounds. Each sampling round involved recording simultaneously the abundance and richness of bees and bee-plant interactions for 15 minutes in the morning and 15 minutes in the afternoon in each patch. The contact of a bee with the reproductive part of a flower was noted as an interaction event. Surveys were conducted on sunny, no-windy days with temperatures above 18ºC. Most bees were identified to genus level in the field, but we collected one individual per morphotype with a sweep net for further species or morphospecies identification in the laboratory (see Cano et al., 2022, 2024, for details). Our sampling patches were consistently placed in diverse floral stands among those available to ensure the detection of the maximum number of bee species and interactions. We aimed to keep their location permanent across sampling rounds, ensuring a minimum separation of 150 m between them whenever possible, although we frequently had to replace them between the two dates of surveys to track floral availability.

While bee abundance and diversity were sampled each year of study, interactions were surveyed after three and five years from the restoration actions (i.e., 2020 and 2022). From these surveys, we obtained the following descriptors: i) Bee abundance: the abundance of bees in patches was obtained as the total sum of bees recorded for each olive farm across patches and sampling rounds for each sampling year; ii) Bee richness: we used the total bee species richness for each olive farm across patches and sampling rounds for each sampling year to estimate species richness from Hill numbers at q=0. Species richness was calculated using the ‘*iNext*’ package in R (Hsieh et al., 2016) with the endpoint doubling the greater bee abundance for each sampling year (minimum species sampling coverage of 0.89); iii) Bee-flower interaction metrics: we employed the plant-bee interaction surveys to build interaction networks from matrices of interaction frequency for each olive farm (interaction pooled across patches and sampling rounds). Then, we calculated the following network metrics: connectance, defined as the number of interactions relative to the maximum possible number, and diversity of interactions (Shannon index). We selected these two metrics because they are established indicators of structural network complexity (Thébault & Fontaine, 2010) and ecosystem function quality (Kaiser-Bunbury & Blüthgen, 2015), respectively. Based on previous studies, we anticipate increased connectance and diversity of interactions in restored networks (Cusser & Goodell, 2013; Forup et al., 2008). Interaction networks and estimation of network metrics were conducted using the ‘*bipartite*’ package in R (Dormann & Strauss, 2014); iv) Pollinator visitation: we used the interaction surveys to obtain the flower visitation rate (FVR) as the number of floral visits by all bees per minute within each olive farm as a proxy of pollination function.

### Predictors associated with the original management intensification of the farm

We considered two categorical predictors of the RD associated with agricultural intensification: ground cover intensification (high-intensity vs. low-intensity) and the agrochemical intensification level (organic vs. non-organic farming). We further considered two continuous predictors, the proportion of herbaceous cover existing before restoration (i.e., 2016) in the olive field and the herb species richness in semi-natural habitat in the same year, the latter as an indicator of potential source of herb species and pollinator spill-over through time, from the semi-natural patches to the floral patches within the olive field (Rey et al., 2019; Tarifa et al., 2021).

### Predictors associated with the heterogeneity at landscape and floral patch scales

We characterised heterogeneity at two scales. At the landscape scale, we considered the landscape complexity within a 1 km radius around the centroid of each olive farm, using descriptors related to the recovery of bee communities and their functions: a) edge density (m/ha) as a landscape configurational metric (Martin et al., 2019), considering as edges the existing land use margins, field margins, boundaries with other farms and crops, and other linear elements such as rivers and streams; b) proportion of non-forest semi-natural habitats, that is, semi-natural areas excluding tree or shrub formations (i.e., floral field margins, and grasslands), considered as favorable open habitats for bees. This last variable is a measure of landscape compositional heterogeneity (Martin et al., 2019). Although we also obtained the proportion of forest and woodland, it was unrelated to any estimator of RD, which could be expected since forests and woodlands in the region are dominated by anemophilous plants (Fagaceae, Pinaceae, Oleaceae and Anacardiaceae). Therefore, we eventually removed it from further analyses. We used recent land-use cartography (data from SIOSE 2016 available at http://www.juntadeandalucia.es/medioambiente/site/rediam/) to estimate these landscape descriptors using a GIS platform (QGIS v.2.14). Extended details may be found in Rey et al., (2019) and Cano et al., (2022).

At the patch scale, we considered the two 10 m^2^ (normally 2 × 5 m) multi-floral patches per farm, where we surveyed bees and their activity. These patches were located within floral stands scattered throughout the olive field in each farm. We recorded floral cover (%) and the occurrence of flowering herb species visually in each patch during every sampling round. Plant species composition within patches differed among localities, reflecting the wide geographic extent of this study.

### Predictors associated with the magnitude of change created by restoration interventions

We considered a series of indicator variables to measure the degree of change in the study farms resulting from the intervention actions. First, we obtained field measurements of the area occupied by entomophilous woody plants within reforested hedgerows successfully established. Then, we expressed these measures relative to farm size to obtain the proportion of the area reforested with entomophilous hedgerows in each olive farm. Second, we employed the following variables that consisted of differences between their post-(in 2020) and pre-restoration interventions values (in 2017): At the farm level, we considered both: a) the herb cover, and b) herb richness within the olive field. To obtain these variables, we established six sampling sites over the total productive area in each farm. In each site, we measured herb richness (within 1 m^2^ square patches) and herb cover (within the area between four olive trees surrounding the 1 m^2^ patches) during April and May of 2017 and 2020 via visual inspection. We further used four 1 m^2^ square patches to estimate herb species richness in non-productive semi-natural habitat patches (woodlands, scrublands, or grasslands, depending on the farm) within or adjacent to each farm. Herb richness and cover in semi-natural or olive field areas were estimated as the mean value between these patches within each olive farm. At the patch level, we considered: a) floral cover and b) floral richness from the floral patches described above, again for the post- (2020) and pre-restoration (2017) intervention periods. Both variables were estimated as the mean value across sampling rounds between the two patches within each olive farm for each period.

### Assessment of the Recovery Debt

Following Moreno-Mateos et al. (2017), we calculated the RD of each target variable using their estimated values at an initial time (*Y*_*s*_) (either immediately before or after restoration actions), the values of that same variable after a certain period had elapsed (*Y*_*e*_), and the reference value (*Y*_*r*_) for each variable from a reference system. Specifically, to produce more reliable comparisons among periods of different lengths, we consistently calculated the RD per year (RD_t_) as follows [Eq. 1]:

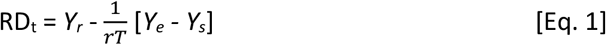

Where *T* is the number of years elapsed between the initial and final estimations of the variables, and *r* is a constant that contains the modelled exponential approximation of the transit from *Y*_*s*_ to *Y*_*e*_ through the expression [Eq. 2]:

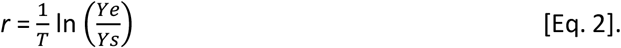

More details of the analytical development involved in the quantification of the RD are provided in Moreno-Mateos et al., (2017). To make the RD_t_ comparable between indicators (abundance, diversity and interaction descriptors), we used a recovery debt ratio by dividing RD_t_ by absolute values of *Y*_*r*_ and expressed it as a percentage.

Thus, we first calculated RD_t_ for the periods 2018-2020 and 2020-2022 to evaluate whether it decreased with time from the implementation of the restoration actions (see Fig. 1). We chose to estimate RD_t_ at two specific time intervals and to evaluate its decrease between them as indicator of RD shortening, rather than comparing the rate of recovery (i.e., *r*, the rhythm or velocity of recovery). We adopted this more conservative approach because estimating a constant rate of increase in diversity based on only three early data points would be less reliable. Afterwards, we calculated the RD_t_ for each target variable from the first data collection (2018 for abundance and richness variables and 2020 for interaction network variables) until the last (2022). We employed these latter values as response variables to assess how they relate to continuous predictors of the original farming intensification and landscape heterogeneity and of ecological contrast generated by restoration interventions.

### Reference system

Assessing RD requires establishing suitable reference systems (Moreno-Mateos et al., 2017), often undisturbed or in a pre-disturbance state (Gann et al. 2019), which is typical in ecological restoration studies (e.g., Dellicour et al., 2023; Devoto et al., 2012; Strobl et al., 2019). Determining a suitable reference system depends on the specific optimal conditions demanded by the group of organisms or ecological function under consideration. However, in some instances, establishing an appropriate reference system can be challenging due to either a scarcity of suitable natural references or their inability to match the desired target state (Harris et al., 2006), particularly in managed systems like agroecosystems. In the specific case of bees in agricultural systems, the ideal reference system should be agricultural landscapes encompassing several features that promote the preservation of bee communities and functions at their best, such as organic farming (Bengtsson et al., 2005; Samnegård et al., 2019), wide cultivation frames allowing a higher proportion of open areas in the cultivated field (Bravo-Monroy et al., 2015), the maintenance of semi-natural areas (Bartual et al., 2019; Cole et al., 2017) and crop diversification at farm and landscape scales (Aguilera et al., 2020; Raderschall et al., 2021). Existing information on olive groves clearly identifies the optimum for the biodiversity of many species groups (Rey et al., 2019; Tarifa et al., 2021), and specifically for bees and the pollination function (Martínez et al., 2019; 2020; Cano et al. 2022; 2025), as farms in relatively complex (heterogeneous) landscapes that are under organic farming and leave herbaceous ground covers. Nonetheless, before estimating RD, its variation, and its relationship with predictors, we validated the suitability of our candidate reference systems for the recovery of bee communities in perennial woody crops such as olive groves. An extensive explanation of the procedure for determining and validating a suitable reference system, further controlling for geographic and spatial autocorrelation among sites of bee species composition, richness and abundance, may be found in Appendix S2. After this validation process, we selected two olive farms as reference systems (labelled “*Ta*” and “*Cas*” in the scatterplots of Fig. S1 and S2; Appendix S2). For each target variable, the reference value was defined as the highest value recorded across these two farms over the three years. For all variables except connectance, this maximum occurred in the final year of the study (2022), indicating that even the reference farms were undergoing recovery following the implemented changes. For this reason, and given the limited number of transformed farms, we retained the reference farms as additional cases in the test sample. Excluding them did not qualitatively affect the results (Appendix S3), either in terms of the overall reduction in recovery debt or in the analysis of the correlates (i.e., landscape simplification, initial ground management intensity, and magnitude of changes triggered by restoration interventions) of the RD remaining at the end of the study.

### Statistical analyses

Non-parametric simple tests were preferable to other alternatives because of the relatively small number of experimental units (i.e., 12 to 14 olive farms subjected to restoration, depending on the bee recovery variable considered). We first evaluated the variation in RD_t_ between 2018-2020 and 2020-2022 using Mann-Whitney tests. We only assessed the variation of RD_t_ between these two periods in abundance and richness, since the bee-plant interaction variables (flower visitation rates and interaction metrics) were not measured in the first period.

Similarly, we used Mann-Whitney tests to compare the RD_t_ of each target variable between olive farms with contrasting herb cover management (high-intensity vs. low-intensity) and with contrasting use of pesticides (non-organic vs organic), considering the whole period elapsed after intervention (that is, from 2016-17 until 2022).

Finally, we assessed through Spearman-rank correlation tests the relations between the RD_t_ for each target recovery variable and the metrics and descriptors of landscape, farming intensification of ground cover, and intensity of change caused by restoration interventions. Prior to these analyses, we checked, through Variance Inflation Factor (VIF) and correlation tests, that no high correlation existed between these descriptors (Appendix S1: Table S3 and Fig. S2, respectively). Then, we conducted separate simple Spearman correlations for each descriptor with RD_t_ due to the limited sample size. Once significant simple correlations were detected, the significant predictors were incorporated into three separate Spearman’s partial correlation analyses with each response variable, each one corresponding to a different group of descriptors: landscape, farming intensification of ground cover, and intensity of change caused by restoration interventions. This allows us to refine relationships with each target RD_t_ response variable while controlling for its possible relationships with other significant predictors.

We conducted Mann-Whitney tests and Spearman’s simple and partial correlations using the *wilcox*.*test, cor* and *pcor*.*test* functions, respectively, from the *stats* and *ppcor* packages in R (R Core Team, 2023). Correlations were visualised using heatmaps generated by the *corrplot* function from the *corrplot* package in R.

## 3- Results

Target variables (bee abundance, richness, flower visitation and interactions metrics) in the reference systems were typically high and relatively stable compared to other farms (i.e., with higher mean and lower coefficient of variation across time; see Appendix S1: Table S4), as expected under optimal or near-optimal conditions for supporting bees and their pollination function in the olive grove landscapes.

### Temporal trend of the recovery debt after restoration interventions

We observed a significant decrease in per-annual RD over time (meaning recovery) for the abundance of bees. That is, RD_t_ was higher in the period from 1st to 3rd year after restoration interventions, and lower for the period elapsed between the 3rd and 5th year after restoration interventions (Mann-Whitney *U*= 32; *p*-value = 0.025; Fig. 2). Similarly, RD_t_ of bee richness decreased over time after restoration interventions although our data provide weak support for this decrease (Mann-Whitney *U*= 54; *p*-value = 0.102; Fig. 2).

**Figure 2.**
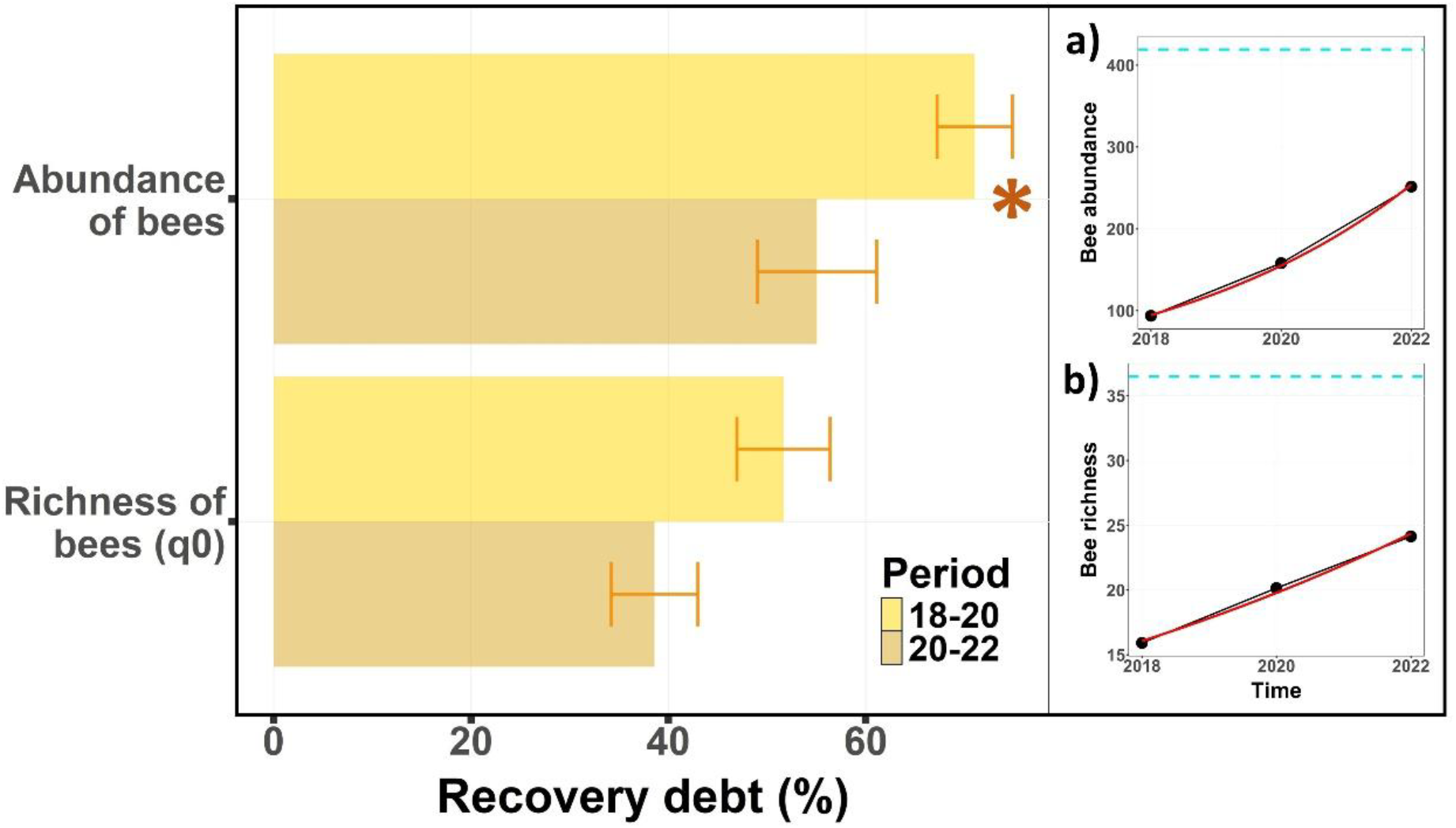
Change between time intervals in the RD of abundance and richness of bees in floral patches. Barplots show mean and standard errors of per-year RD (i.e., RD_t_) of abundance (i.e., abundance of bees in floral patches) and richness for bee assemblages in floral patches during the two periods evaluated (i.e., periods from 2018-2020 and 2020-2022). Asterisks show significant differences between periods. Note that the RD_t_ decreases between consecutive time intervals from the restoration interventions. Here and in the next Figures, a larger RD_t_ means that the ecological attribute is further from (or has a larger deficit compared to) the reference system. The right subpanel shows mean raw values (dots connected by a black line) across farms for each time period for (a) bee abundance and (b) bee richness, along with their exponential growth trend (red line) of approaching the reference system value (dashed blue line). Comparison of barplot and trends in the right panels allows linking increasing mean values in the community attribute to decreasing RD_t_ over time.

### Effects of the pre-intervention level of farming intensity on the recovery debt

Neither the intensity of the herb cover management nor pesticide use showed any significant effect on RD_t_ of any target variable (Appendix S1: Table S5; Fig. 3a and 3b, respectively). We observed a tendency (but no significant difference) for lower RD_t_ of the richness of bees in organic farms (Fig. 3b) and for connectance in high-intensity farms (Fig. 3a).

**Figure 3.**
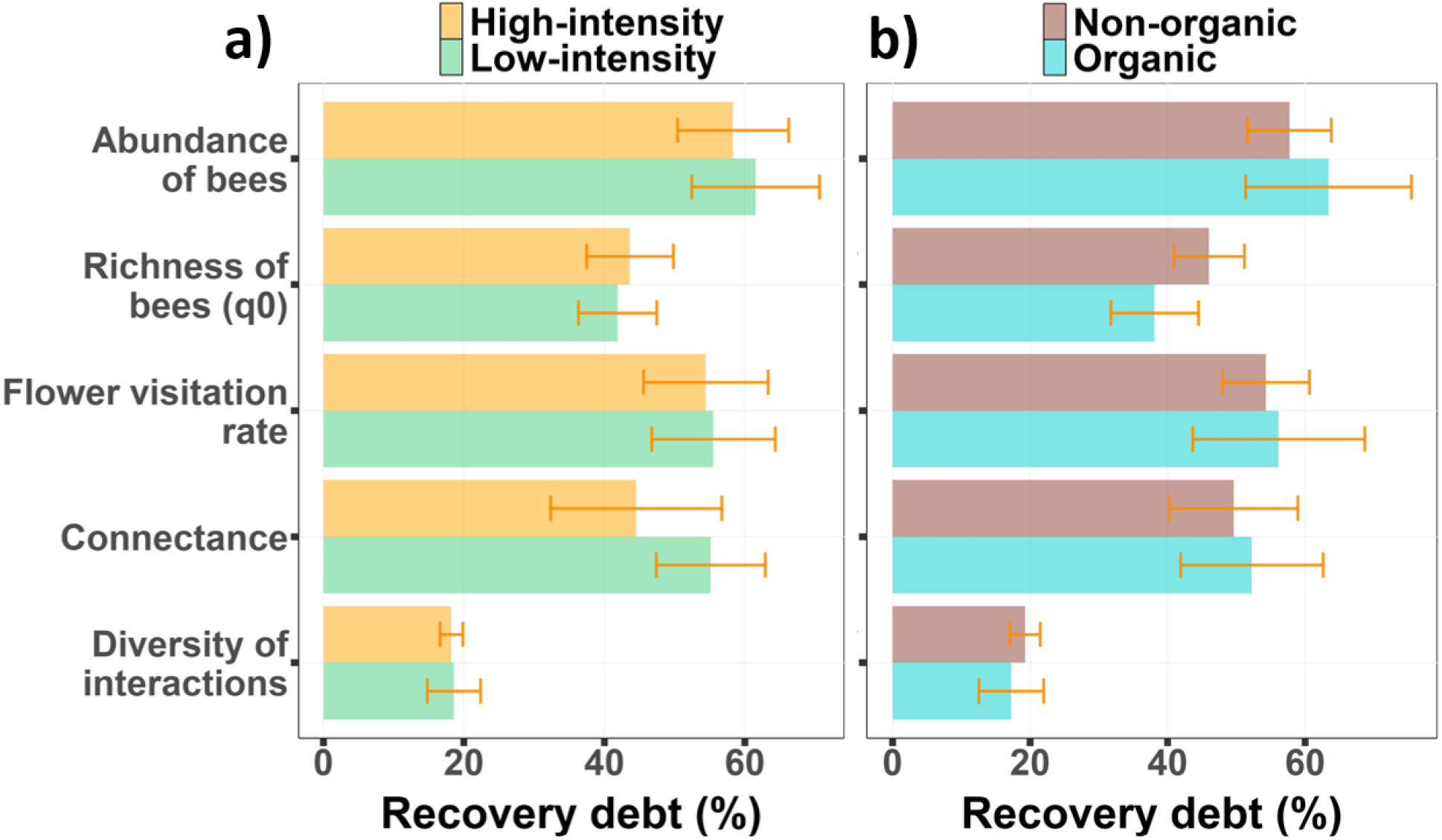
Recovery debt as a function of pre-intervention management intensity. Barplots show means and standard errors of per-year RD (i.e., RD_t_) for bee abundance and richness in floral patches and plant-bee network metrics (i.e., flower visitation rate, connectance and diversity of interactions) for a) each pre-intervention herb cover management type and b) type of pesticide use. In this case, RD_t_ for each target variable (expressed in relative terms, (%), was calculated from the first data collection (2018 for abundance and richness variables and 2020 for interaction network variables) until the last year of monitoring (2022). None of the comparisons was statistically significant at *p* < 0.05 level. Note that a larger recovery debt means that the ecological attribute is further from the reference system.

### Pre-intervention ground cover and landscape heterogeneity as predictors of recovery debt

We found a significant decrease in RD_t_ of the flower visitation rate associated with the proportion of semi-natural non-forest cover (r = -0.58; *p*-value = 0.052; Fig. 4). Likewise, higher herb richness in the semi-natural areas of the olive farms was associated with a significant decrease in RD_t_ of the flower visitation rate (r = -0.73; *p*-value = 0.007; Fig. 4), probably because it also entailed recovery of abundance of bees (r = -0.59; *p*-value = 0.027; Fig. 4).

**Figure 4.**
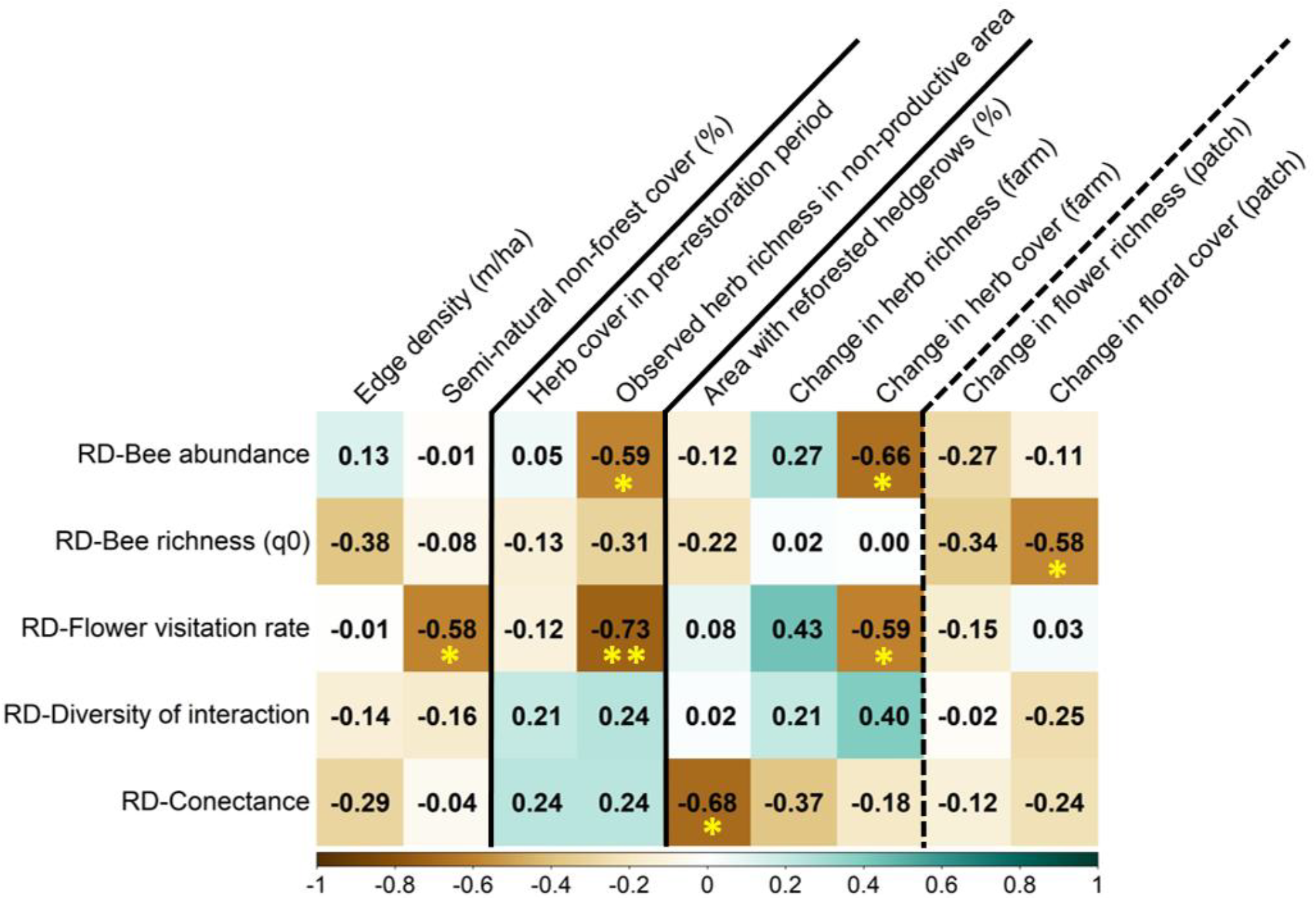
Spearman’s correlations between biodiversity RD and environmental variables to show which pre-intervention conditions and intervention efforts favor less recovery debt (i.e., shortening of RD_t_). The heatmap shows the degree of correlation between the per-year RD (i.e., RD_t_) of target variables (rows) and, from left to right, the landscape simplification descriptors (columns 1 and 2), the predictors associated to the original management intensification of the farm (columns 3 and 4) and the magnitude of change triggered by restoration interventions at farm and floral patch level (columns 5-7 and 8-9, respectively). RD_t_ for each target variable was calculated from the first data collection (2018 for abundance and richness variables; 2020 for interaction network variables) until the last year of monitoring (2022). Note that negative correlations identify variables (pre-intervention conditions and magnitude of changes after intervention) that led to a smaller deficit in the specific pollinating insects and function attribute or shorter time lag in paying off the recovery debt. Asterisks show significant Spearman’s correlation tests (* *p* < 0.05; ** *p* < 0.01).

### Change in olive farms triggered by restoration interventions and the recovery debt

We observed a significant decrease in the RD_t_ of bee abundance and flower visitation rate with the change between the pre- and post-restoration periods in herb cover at the farm scale (respectively: r = -0.66, *p*-value = 0.012 and r = -0.59, *p*-value = 0.040; Fig. 4). The RD_t_ of bee species richness significantly decreased as floral cover difference in the floral patches increased between the pre- and post-restoration periods (r = -0.58; *p*-value = 0.033; Fig. 4). Finally, the RD_t_ of connectance of bee-plant networks significantly decreased with a higher proportion of area successfully reforested with hedgerows (r = -0.68; *p*-value = 0.025; Fig. 4).

### Combined effect of pre-intervention state and change caused by interventions in the RD

All significant associations of each RD_t_ estimator with pre-intervention states (farming intensification and landscape heterogeneity) and the changes caused by restoration interventions (i.e., significant simple correlations) were maintained and even strengthened after incorporating them into Spearman’s partial correlations (Appendix S1: Table S6).

## 4- Discussion

It must be stressed that our results reflect the incipient process of recovery (5 years) after the implementation of restoration actions. In spite of this limited time, we found that implementing restoration measures in olive crops already shortens the recovery debt of bee abundance in the short term. The level of shortening of the RD_t_ was however not related to the farming practices (intensity of ground management or pesticide application) existing before the intervention, but it seems to be more related to the quantity and quality of the non-forest semi-natural habitat in the landscape and to the ecological contrast generated by the interventions. As expected, they influenced the bee abundance and flower visitation rate to a larger extent than the bee richness and interaction network properties, suggesting that these more complex descriptors of the community take longer to recover after active restoration or application of agri-environmental measures in agricultural fields. Our findings are thus aligned with other studies highlighting that the maintenance and early recovery of an efficient pollination service may be more effective and easier, focusing on pollinator abundance and pollination rates (Genung et al., 2017; Winfree et al., 2018), rather than directly attempting the recovery of interaction network properties (but see Kaiser-Bunbury et al., 2017). Some conclusions of this study should be interpreted with caution due to the limited statistical power arising from the small number of farms in our test sample. However, this work can be considered a proof of concept illustrating the usefulness of the Recovery Debt approach for evaluating the outcomes of active restoration practices in general, and for assessing biodiversity recovery in agricultural landscapes following the implementation of agri-environmental measures and schemes in particular.

### Short-term recovery of bee abundance and richness after restoration interventions

The significant decrease in the RD_t_ of bee abundance (meaning recovery) observed over time indicates that bee abundance quickly responds to restoration measures. Pollinating insects, including bees, have been consistently observed to respond rapidly to environmental disturbances (Kuussaari et al., 2009; Sexton & Emery, 2020) and, as this study shows, this also seems to be the case after implementation of active restoration measures. Additionally, it is commonly observed that years with high floral resource availability lead to an increase in the population size of the next generations of bees (Cusser et al., 2015; Williams & Kremen, 2007), reasonably explaining the observed pattern in olive farms after interventions promoting ground herb covers.

Bee richness in floral patches followed the same trend of bee abundance, but the decrease of RD_t_ over time was not significant. Unlike abundance, which responds rapidly to variations in floral resources and thus allows an earlier detection of changes (Crone, 2013; Cusser et al., 2015), recovery of richness may take longer to become evident. Consequently, even five years after restoration in olive farms, the RD_t_ of bee richness did not consistently decrease over time.

It is noteworthy that the values obtained for the RD_t_ of bee abundance and richness fall within the range observed in agroecosystems over longer periods (i.e., 20-30 years; Moreno-Mateos et al., (2017). This suggests that the restoration measures implemented in the olive farms, primarily through changes in herb cover management, are highly effective in restoring bee abundance and, to a lesser extent, richness.

### Effects of pre-intervention farming intensity on the recovery debt

We did not find evidence of lower RD_t_ in any bee community or bee-plant interaction descriptors in farms that had less intensive pre-intervention farming, either based on ground herb cover management or in pesticide application. This contradicts studies in other systems showing high responsiveness of pollinators to organic farming (Bloom et al., 2023; Holzschuh et al., 2007; Kennedy et al., 2013) and grassland-type restorations (Kaiser-Bunbury et al., 2017; Lane et al., 2020; Noreika et al., 2019; Traveset et al., 2024). In fact, previous studies conducted before interventions in the same studied olive farms have shown that low-intensity herb cover management and organic farming have benefits for above-ground cavity-nesting bees (Martínez-Núñez et al., 2020a) and bee-plant interaction networks (Cano et al., 2024; Martínez-Núñez et al., 2019), so it would be reasonable to expect lower recovery debt under these circumstances. Part of this absence of effects may be due to a lack of statistical power for detecting significant effects because of a limited number of farms subjected to restoration and with repeated monitoring of bees after the interventions, which is an inherent common challenge in these types of studies. In fact, there were some trends, despite not being significant, that point to decreased RD_t_ in the diversity of bees and the diversity of interactions, mainly under organic farming. This trend should be confirmed through longer term monitoring. In any case, we can discuss some possible alternative explanations. Some studies have remarked that restoration strategies focusing on key plant species generate a faster response from pollinators than just focusing on overall plant and flowering abundance and diversity (Sexton & Emery, 2020). In fact, Martínez-Núñez et al., (2020b) found that some flowering species may be fundamental in promoting a core of interactions with cavity-nesting bees in olive groves under any management. In any case, the lack of effect of the pre-intervention intensification levels on the existing RD_t_ may alternatively be explained by the moderator effects of the landscape context of each farm, offering habitats that act as a source of bees for the studied floral patches in the olive field through spillover (Tscharstke et al. 2005; Schepper et al. 2013), thus buffering the effect of the intensification practices. Likewise, the RD_t_ might be more related to the ecological contrast, in terms of enhancement of floral and other resources for bees, caused by the level of interventions (Schepper et al. 2013; 2015; Marja et al. 2019). These effects are discussed below.

### The quality of semi-natural patches and the landscape context to some extent determine the RD of bees and its function in floral patches within the olive field

The analyses of the effects of the landscape context on the RD_t_ revealed that semi-natural areas with higher herb richness favoured the recovery of both the bee abundance and the FVR in floral patches within the olive crop matrix. Additionally, the positive impact of non-forest semi-natural areas on the FVR indicates that these areas provide substantial support to bees and their functions when they have high-quality floral resources (e.g., Bartual et al., 2019; Cano et al., 2022; Dellapiana et al., 2025; Larkin & Stanley, 2023), thus explaining the increase of bee abundance in the floral patches of the olive farms through its spillover from the semi-natural areas (Garibaldi et al., 2011; Ponisio et al., 2019; Ragué et al., 2022). The observed rise in the FVR, suggests a potential enhancement in the pollination service for plants in the floral patches scattered through the olive farms’ ground cover. Therefore, the presence and maintenance of high-quality semi-natural areas with abundant floral resources may play a role in recovering bee populations and improving pollination services within agricultural landscapes. Our results thus support the moderator effect that the surrounding landscape context (particularly suitable semi-natural habitat) has on the impact of intensive agriculture on bees and their pollination function (Schepper et al. 2013; 2015; see also Cano et al. 2025, for the case of functional diversity of bees in olive groves). Moreover, here we demonstrate that this moderator role of suitable habitat for bees works not only for determining the abundance and function of bees and their RD_t_, but also for recovery, as shown by a significant reduction of RD_t_ of bee abundance (and marginally in richness) through time intervals after interventions.

### The ecological contrast promoted by the intervention aimed to increase floral covers in olive farms also determines the recovery debt

Increasing the herb cover at the farm level over time had the greatest impact on the recovery of bee populations and their function, as shown by its negative relationship with the RD_t_ of bee abundance and flower visitation rates (FVR). This increase in herb cover was the consequence of shifting the intensity of herb cover management from aggressive farming practices, which leave the soils bare through herbicide application and/or deep tillage, to a management consisting of leaving herbaceous cover most of the year and mechanically removing it through mowing at late spring when many flowering plants have already completed their life cycles. This type of low-intensity management is a fundamental element of the eco-regimes approved in Andalusia for olive groves and other permanent croplands within the new European Common Agricultural Policy (Pérez-Pozueto et al., 2024), which focuses on the maintenance of herbaceous cover, at least in the inter-rows of olive trees. Its positive effects on many different groups of organisms are widely acknowledged through comparative studies (Rey et al., 2019; Tarifa et al., 2021), including pollinating insects in olive groves (Cano et al., 2022; Martínez-Núñez et al., 2020a). Our study provides experimental evidence in olive groves that changes in the ground herb cover at the farm scale are associated, over time, with increases in bee abundance (i.e., since it was negatively correlated with RD across sites) as the available floral resources become more abundant (see also Cusser et al., 2015; Williams & Kremen, 2007 in other systems). The resulting increase in bee abundance in floral patches would, in turn, enhance the FVR, thereby ensuring the pollination service in the floral patches (Garibaldi et al., 2011; Groom, 1998). Moreover, the extent of change in herbaceous or floral cover more frequently relates, across farms, to the RD_t_ of bee abundance, richness, and FVR, confirming the relevant role of the ecological contrast created by restoration or application of agri-environmental measures (Marja et al. 2019). Furthermore, since no relationship was found between RD_t_ and the type of herb cover management in the pre-restoration period, these results suggest that the degree of change generated by the restoration actions is more important than the pre-intervention state to determine the level of recovery achieved in bees and the pollinator function. Our results are in line with conceptual proposals and observational studies that remark that, in the case of pollinating insects, the ecological contrast promoted by agri-environmental measures, rather than the farming type, is key for bees’ recovery (Marja et al., 2019; Scheper et al., 2013).

Interestingly, the scale at which the ecological contrast caused by interventions works to recover bees was different when focusing on the abundance or bee diversity. We detected a reduction in RD_t_ (meaning a recovery) of bee species richness with the enhancement of floral cover in the floral patch scale but not at the farm scale. This is consistent with other studies showing that pollinator richness responds better to the availability of floral resources at smaller scales (e.g., Hegland & Boeke, 2006), explaining their positive response to the ecological contrast triggered at the floral patch level. This remarks the need to warrant that our interventions enhance the herbaceous and floral cover at both farm and patch scales (Scheper et al., 2015).

Reforested hedgerows only enhanced bee-plant network complexity in terms of connectance, with less RD_t_ corresponding to a greater proportion of area of restored hedgerows in olive farms (i.e., negative correlation). Although increased network connectance can result from a reduction in the number of interacting species (Soares et al., 2017), this does not seem to be the case here, as the proportion of area restored with hedgerows did not affect the RD_t_ of bee richness. Instead, greater landscape connectivity through a higher proportion of green infrastructures is known to shape bee community composition (Kremen et al., 2018; Öckinger et al., 2018; Traveset et al., 2024), and it is an expected result in restored networks (Cusser & Goodell, 2013; Forup et al., 2008). In the olive groves of this study, such connectivity could be further enhanced because many of the planted woody species have entomophilous flowers.

This pattern is consistent with Ponisio et al. (2019), who found that restored hedgerows promote long-term persistence and support pollinator metacommunity dynamics in intensively managed agricultural landscapes. Thus, olive farms with more restored hedgerows may have altered community composition by favouring more generalistic bees from semi-natural areas, as documented in Cusser & Goodell (2013), increasing the number of unique interactions and therefore leading to higher connectance. However, further studies are needed to confirm this.

## 5- Conclusions

Our study demonstrates that active ecological restoration, through wildlife-friendly herb cover management in olive farms, can effectively restore bee communities in the short term. The extent of this recovery largely depends on the surrounding landscape context, and on the ecological contrast created by restoration actions, which enhance bee abundance, richness and function. This suggests that restoration success in agricultural systems depends not only on local practices but also on broader spatial planning. Meanwhile, the poor response of interaction-network properties underscores the need for long-term commitments to fully re-establish ecological complexity and resilience.

Beyond documenting effective recovery, our study highlights the value of the recovery debt framework as a prospective tool to model and monitor biodiversity restoration through time after restoration interventions in agricultural systems. While long-term monitoring remains essential to accurately estimate recovery trajectories and rates, these results represent a significant initial step toward understanding how active ecological restoration contributes to recovering pollinator community complexity in permanent agroecosystems and in guiding effective restoration and adaptive management plans.

## Acknowledgements

We thank the owners of the olive orchards who granted us access to their properties. This work is part of the projects RECOVECOS (PID2019-108332GB-I00, funded by MICIN/AEI/10.13039/501100011033) and OLIVARES VIVOS + (LIFE20 AT/ES/001487, European Commission). Restoration and other interventions in the olive fields were funded by the LIFE project OLIVARES VIVOS (LIFE14 NAT/ES/1001094). We also thank Antonio López-Orta for support in assessing restored hedge dimensions and Ana González Robles, Gemma Calvo and Francisco Camacho for assistance in herb cover and richness determination. Participation of AJP in bee and flower censuses and data curation was funded by MICINN through the European Regional Development Fund [SUMHAL, LIFEWATCH-2019-09-CSIC-4, POPE 2014-2020]. D.C. was granted a predoctoral fellowship (FPU17/02186). CMN was supported by a “Juan de la Cierva” postdoctoral grant (ref. FJC2021-046829-I) and by a Junta de Andalucía (ref. DGP_POST_2024_00228) Postdoctoral Fellowship, funded by the Spanish MICINN, and the Andalusian Regional Government.

## Authors’ contributions

P.J.R. conceived the ideas and designed the methodology. D.C. and A.J.P. conducted the bee surveys and identified the collected bees. A.J.P. and R.T. conducted monitoring of herb cover communities and assessed the dimensions of the restored hedges. T.S. processed the land use cartography data and produced metrics of the landscape heterogeneity. C.R. and J.E.G provided logistic support to field work and planned, executed and supervised restorations and other interventions in the olive farms. D.C, P.J.R. and J.M.A. analysed the data. D.C. and P.J.R led the writing of the manuscript with input from C.M-N and J.M.A.

## Notes

https://doi.org/10.6084/m9.figshare.30609599

### Competing Interest Statement

The authors have declared no competing interest.

