## Appendix S1 for "Assessing pollinator community recovery in restored agroecosystems using the recovery debt framework"

**Affiliations:**

**Appendix S1**

**Table S1.-** Detailed information about the farming management and landscape characteristics of the study olive farms.

| Locality | Acronym | Original herb cover management | Use of pesticide | Farm size | % semi-natural cover in 1 km radius buffer | % forest cover in 1 km radius buffer | % no-forest semi-natural cover in 1 km radius buffer | % olive tree cover in 1 km radius buffer | Richness of land use/cover in 1 km radius buffer | Shannon diversity of total land use/cover in a 1 km radius buffer | Largest patch index (%) | Edge density (m/ha) | Mean patch area (ha) | Distance to nearest patch of same land use (m) |
| --- | --- | --- | --- | --- | --- | --- | --- | --- | --- | --- | --- | --- | --- | --- |
| Torredonjimeno | Be | High-intensity | No organic | Small | 2.297 | 0.995 | 1.302 | 97.162 | 10 | 0.182 | 50.153 | 91.728 | 4.984 | 210.170 |
| Castro del Río | Ca | High-intensity | No organic | Small | 1.880 | 1.205 | 0.675 | 76.502 | 11 | 0.767 | 19.512 | 117.927 | 2.046 | 57.208 |
| Peal de Becerro | Cg | Low-intensity | No organic | Large | 14.104 | 12.882 | 1.222 | 83.978 | 11 | 0.615 | 33.866 | 85.379 | 5.150 | 136.836 |
| Lantejuela | Dl | Low-intensity | Organic | Small | 5.033 | 4.675 | 0.358 | 64.165 | 11 | 0.947 | 30.940 | 121.518 | 2.533 | 70.778 |
| Pegalajar | Cas | Low-intensity | Organic | Large | 60.174 | 42.368 | 17.806 | 36.744 | 14 | 1.378 | 19.212 | 196.877 | 1.593 | 65.236 |
| Mancha Real | Mon | Low-intensity | No organic | Large | 22.051 | 21.743 | 0.308 | 72.743 | 11 | 0.824 | 51.279 | 98.739 | 6.059 | 89.823 |
| Moraleda de Zafayona | Mo | High-intensity | No organic | Small | 6.000 | 2.173 | 3.826 | 51.486 | 11 | 1.258 | 16.809 | 184.248 | 3.090 | 127.238 |
| Obejo | Olu | Low-intensity | Organic | Large | 9.339 | 8.918 | 0.421 | 87.962 | 8 | 0.492 | 26.749 | 187.177 | 1.507 | 69.767 |
| Larva | Oli | Low-intensity | Organic | Small | 48.345 | 20.070 | 28.275 | 43.322 | 15 | 1.680 | 11.787 | 211.395 | 2.575 | 80.366 |
| Coripe | Pg | High-intensity | No organic | Small | 13.504 | 13.504 | 0.000 | 43.579 | 15 | 1.436 | 33.990 | 199.202 | 0.618 | 37.167 |
| Santiago de Calatrava | Pc | High-intensity | No organic | Large | 5.066 | 2.537 | 2.529 | 79.373 | 11 | 0.723 | 35.187 | 96.112 | 3.678 | 157.560 |
| Alcaudete | To | Low-intensity | Organic | Large | 3.190 | 3.165 | 0.025 | 93.242 | 11 | 0.368 | 60.142 | 123.401 | 5.421 | 143.531 |
| Nueva Carteya | Toq | High-intensity | No organic | Large | 5.579 | 5.555 | 0.023 | 93.756 | 7 | 0.299 | 74.529 | 83.806 | 4.013 | 48.094 |
| Torres de Albanchez | Ta | Low-intensity | Organic | Small | 31.849 | 28.196 | 3.653 | 59.542 | 11 | 1.240 | 15.046 | 169.616 | 2.377 | 103.944 |

**Table S2.-** Detailed information about the magnitude of change provoked by the recovery interventions implemented in the study olive farms.

| Locality | Acronym | Length of recovered herbaceous boundaries (m) | Total area of intervention on herbaceous cover (m^2^) | % farm area of intervention on herbaceous covers | Area of reforestation in areal stands (m^2^) | Total length of reforested hedges (m) | % farm area initially reforested | % farm area covered by successful reforestation | % farm area initially reforested with hedgerows | % farm area successfully reforested with hedgerows | Number of planted woody plant species | Number of nesting or refuge infrastructures for birds | Area of water ponds installed or restored (m^2^) | Number of installed artificial nest boxes for above ground cavity nesting bees |
| --- | --- | --- | --- | --- | --- | --- | --- | --- | --- | --- | --- | --- | --- | --- |
| Torredonjimeno | Be | 690 | 26108 | 22.425 | 0 | 2437 | 9.259 | 5.675 | 9.145 | 5.605 | 33 | 14 | 7 | 4 |
| Castro del Río | Ca | 0 | 154006 | 88.351 | 0 | 3315 | 3.081 | 0.363 | 3.055 | 0.360 | 27 | 5 | 7 | 4 |
| Peal de Becerro | Cg | 0 | 1015 | 0.032 | 4057 | 7432 | 0.981 | 0.068 | 0.545 | 0.038 | 41 | 18 | 21 | 6 |
| Lantejuela | Dl | 976 | 2329 | 0.113 | 0 | 4904 | 0.806 | 0.159 | 0.238 | 0.047 | 31 | 13 | 7 | 4 |
| Pegalajar | Cas | 0 | 368 | 0.057 | 1161 | 2252 | 1.076 | 0.112 | 0.603 | 0.063 | 17 | 7 | 7 | 6 |
| Mancha Real | Mon | 0 | 624 | 0.017 | 3572 | 4357 | 0.337 | 0.061 | 0.154 | 0.028 | 33 | 30 | 7 | 6 |
| Moraleda de Zafayona | Mo | 1056 | 29550 | 23.127 | 0 | 2896 | 2.877 | 1.033 | 2.876 | 1.033 | 30 | 11 | 0 | 4 |
| Obejo | Olu | 0 | 680 | 0.137 | 0 | 6226 | 1.531 | 0.252 | 1.477 | 0.243 | 33 | 21 | 7 | 6 |
| Larva | Oli | 0 | 990 | 0.461 | 237 | 1789 | 2.294 | 0.599 | 1.896 | 0.495 | 25 | 6 | 7 | 4 |
| Coripe | Pg | 0 | 56477 | 57.017 | 0 | 1368 | 2.383 | 0.771 | 2.255 | 0.730 | 26 | 2 | 0 | 4 |
| Santiago de Calatrava | Pc | 1313 | 660699 | 88.630 | 0 | 10777 | 1.592 | 0.173 | 1.577 | 0.171 | 28 | 13 | 7 | 6 |
| Alcaudete | To | 0 | 0 | 0.000 | 15303 | 3238 | 1.637 | 0.076 | 0.267 | 0.012 | 34 | 12 | 7 | 6 |
| Nueva Carteya | Toq | 1368 | 3982 | 0.495 | 843 | 8964 | 2.364 | 0.774 | 2.099 | 0.687 | 42 | 10 | 0 | 5 |
| Torres de Albanchez | Ta | 0 | 2062 | 0.711 | 1560 | 2209 | 1.615 | 0.339 | 0.898 | 0.188 | 39 | 12 | 0 | 6 |

**Table S3.-** Variance inflation factor (VIF) values of each explanatory variable (i.e., descriptors of landscape simplification and magnitude of change provoked by the recovery measurements implemented in the study olive farms) for each response variable (RD associated to each target variable). The VIF reports how much the variance of the estimated coefficients is inflated when multicollinearity exists. VIF is the inverse of tolerance (1-*R*^2^), where *R*^2^ is the unadjusted coefficient of determination for regressing each independent variable against the others.

|  | **Edge density (m/ha)** | **Semi-natural non-forest cover (%)** | **Pre-operational herb cover** | **Observed herb richness in unproductive area** | **Area with reforested hedgerows (%)** | **Difference for herb richness (farm)** | **Difference for herb cover (farm)** | **Difference for flower richness (patch)** | **Difference for floral cover (patch)** |
| --- | --- | --- | --- | --- | --- | --- | --- | --- | --- |
| **a) RD-Abundance of bees in floral patches** | | | |  |  |  |  |  |  |
| VIF | 1.51 | 1.51 | 1.24 | 1.24 | 2.34 | 1.33 | 2.47 | 2.68 | 2.32 |
| Tolerance | 0.66 | 0.66 | 0.81 | 0.81 | 0.43 | 0.75 | 0.40 | 0.37 | 0.43 |
| **b) RD-Species richness (q=0)** | | |  |  |  |  |  |  |  |
| VIF | 1.51 | 1.51 | 1.24 | 1.24 | 2.34 | 1.33 | 2.47 | 2.68 | 2.32 |
| Tolerance | 0.66 | 0.66 | 0.81 | 0.81 | 0.43 | 0.75 | 0.40 | 0.37 | 0.43 |
| **c) RD-Flower visitation rate** | | |  |  |  |  |  |  |  |
| VIF | 1.26 | 1.26 | 1.38 | 1.38 | 2.92 | 1.76 | 2.13 | 1.47 | 1.44 |
| Tolerance | 0.80 | 0.80 | 0.72 | 0.72 | 0.34 | 0.57 | 0.47 | 0.68 | 0.69 |
| **d) RD-Connectance** | |  |  |  |  |  |  |  |  |
| VIF | 1.26 | 1.26 | 1.38 | 1.38 | 2.92 | 1.76 | 2.13 | 1.47 | 1.44 |
| Tolerance | 0.80 | 0.80 | 0.72 | 0.72 | 0.34 | 0.57 | 0.47 | 0.68 | 0.69 |
| **e) RD-Diversity of interactions** | | |  |  |  |  |  |  |  |
| VIF | 1.26 | 1.26 | 1.38 | 1.38 | 2.92 | 1.76 | 2.13 | 1.47 | 1.44 |
| Tolerance | 0.80 | 0.80 | 0.72 | 0.72 | 0.34 | 0.57 | 0.47 | 0.68 | 0.69 |

**Table S4.-** Mean values and coefficients of variation for each target variable, calculated over the three-year period. Bold values represent the two highest mean values for each target variable across farms, and green-highlighted cells indicate the olive farm that serves as the reference system for each target variable.

|  | **Bee abundance** | | **Bee richness** | | **Diversity of interactions** | | **Connectance** | | **Flower visitation rate** | |
| --- | --- | --- | --- | --- | --- | --- | --- | --- | --- | --- |
| **Farm_code** | **Mean** | **Coefficient of variation** | **Mean** | **Coefficient of variation** | **Mean** | **Coefficient of variation** | **Mean** | **Coefficient of variation** | **Mean** | **Coefficient of variation** |
| Be_E | 205.50 | 100.13 | 14.49 | 53.72 | **_** | **_** | **_** | **_** | **_** | **_** |
| Ca_E | 135.00 | 17.81 | 15.90 | 46.60 | 2.63 | 4.94 | 0.17 | 34.94 | 1.13 | 17.81 |
| **Cas_E** | 135.33 | 49.13 | 27.63 | 24.34 | **3.08** | 5.85 | 0.12 | 24.96 | 1.38 | 34.08 |
| Cg_E | 175.33 | 48.83 | 12.83 | 11.05 | 2.25 | 3.11 | **0.20** | 3.63 | 1.87 | 9.47 |
| Dl_E | 83.00 | 83.49 | 24.39 | 1.49 | 2.72 | 12.60 | 0.16 | 25.87 | 0.69 | 83.49 |
| Olu_E | 157.50 | 90.24 | 17.68 | 36.46 | 2.86 | 2.38 | 0.18 | 57.76 | 1.31 | 90.24 |
| Mon_E | 180.67 | 54.81 | 20.81 | 18.05 | 2.76 | 0.39 | **0.21** | 34.36 | 1.52 | 76.93 |
| Mo_E | **509.00** | 33.62 | **29.07** | 9.29 | 2.60 | 3.41 | 0.15 | 27.31 | **4.24** | 33.62 |
| Oli_E | 27.00 | 10.48 | 14.20 | 8.56 | **_** | **_** | **_** | **_** | **_** | **_** |
| Pg_E | 156.00 | 38.98 | 23.98 | 46.36 | 2.45 | 10.75 | 0.14 | 5.70 | 1.30 | 38.98 |
| Pc_E | 173.67 | 74.46 | 20.46 | 41.02 | 2.88 | 21.42 | 0.18 | 74.41 | 1.84 | 63.99 |
| To_E | 180.00 | 12.58 | 21.65 | 16.29 | 2.36 | 46.44 | 0.18 | 36.66 | 1.41 | 11.26 |
| Toq_E | 107.67 | 44.59 | 19.70 | 46.28 | **2.98** | 25.16 | 0.14 | 6.78 | 1.12 | 15.83 |
| **Ta_E** | **383.00** | 13.29 | **29.23** | 35.01 | 2.37 | 21.23 | 0.13 | 55.29 | **3.19** | 13.29 |

**Table S5.-** Results from the Mann-Whitney test comparing the per year RD (RD_t_) for the sets of abundance, diversity and interaction network variables between categories of original (pre-intervention) herb cover management and use of pesticides in the olive farms. Statistical significance was determined at p < 0.05. RD_t_ for each target variable was calculated from the first data collection (2018 for abundance and richness variables and 2020 for interaction network variables) until the last (2022). Larger RD values means higher deficits in the ecological attribute compared to the refernce system.

| **Target variable** | **Category of agricultural intensification** | **Factor** | **Mean ± SE** | **Median** | ***U*** | ***P*-value** |
| --- | --- | --- | --- | --- | --- | --- |
| Abundance of bees in floral patches | Herb cover management | High-intensity | 58.303 ± 7.90 | 62.207 | 20 | 0.662 |
|  |  | Low-intensity | 61.514 ± 9.08 | 66.956 |  |  |
|  | Use of pesticides | Non-organic | 57.708 ± 6.08 | 62.207 | 16 | 0.344 |
|  |  | Organic | 63.378 ± 12.04 | 68.729 |  |  |
| Bee assemblages´ species richness | Herb cover management | High-intensity | 43.623 ± 6.18 | 42.187 | 25 | 0.949 |
|  |  | Low-intensity | 41.867 ± 5.57 | 39.478 |  |  |
|  | Use of pesticides | Non-organic | 46.024 ± 5.11 | 43.006 | 33 | 0.282 |
|  |  | Organic | 38.082 ± 6.37 | 34.150 |  |  |
| Flower visitation rate | Herb cover management | High-intensity | 54.415 ± 8.87 | 63.731 | 18 | 1.000 |
|  |  | Low-intensity | 55.501 ± 8.78 | 61.160 |  |  |
|  | Use of pesticides | Non-organic | 54.277 ± 6.33 | 61.250 | 15 | 0.755 |
|  |  | Organic | 56.128 ± 12.50 | 61.160 |  |  |
| Connectance | Herb cover management | High-intensity | 44.499 ± 12.18 | 26.812 | 11 | 0.343 |
|  |  | Low-intensity | 55.109 ± 7.77 | 51.304 |  |  |
|  | Use of pesticides | Non-organic | 49.592 ± 9.34 | 51.304 | 14 | 0.638 |
|  |  | Organic | 52.222 ± 10.36 | 38.034 |  |  |
| Diversity of interactions | Herb cover management | High-intensity | 18.197 ± 1.62 | 18.050 | 19 | 0.876 |
|  |  | Low-intensity | 18.556 ± 3.79 | 15.498 |  |  |
|  | Use of pesticides | Non-organic | 19.248 ± 2.18 | 18.050 | 21 | 0.638 |
|  |  | Organic | 17.228 ± 4.71 | 15.498 |  |  |

**Table S6**. Table of Spearman partial correlations of RD with the predictors of pre-intervention state and change after intervention, which kept simple significant correlations with the target RD variable. Asterisks show significant correlations (i.e., (*) 0,01 < *p*-value < 0,05; (**) 0,001 < *p-value* < 0,01).

|  | **Semi-natural non-forest cover (%)** | **Observed herb richness in non-productive area** | **Area with reforested hedgerows (%)** | **Change in herb cover (farm)** | **Change in floral cover (patch)** |
| --- | --- | --- | --- | --- | --- |
| **RD-Bee abundance** |  | -0.64 * |  | -0.70 ** |  |
| **RD-Bee richness (q0)** |  |  |  |  | -0.58 * |
| **RD-Flower visitation rate** | -0.62 * | -0.74 * |  | -0.79 ** |  |
| **RD-Conectance** |  |  | -0.68 * |  |  |

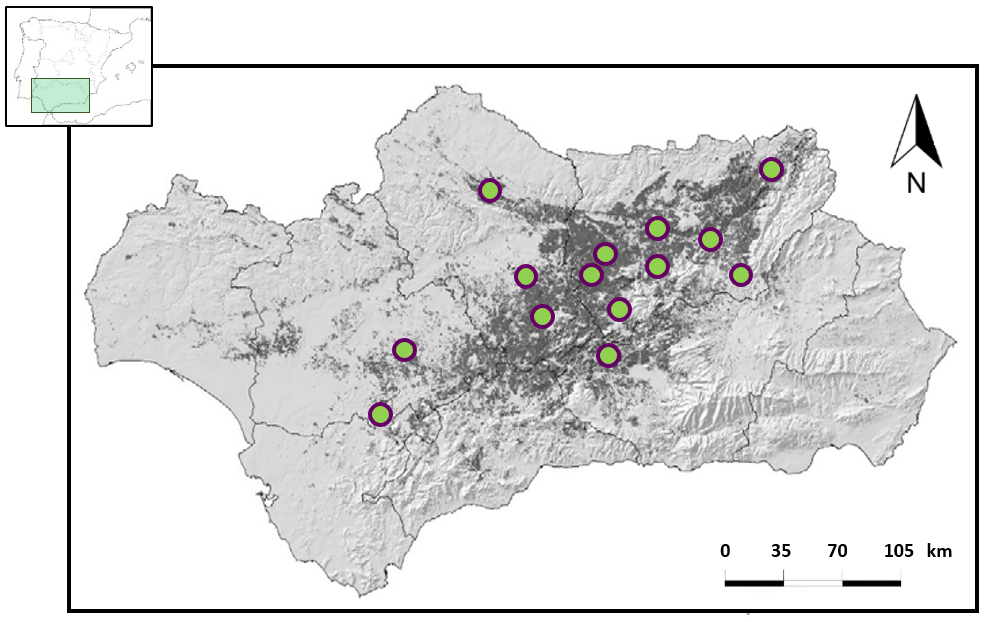

**Figure S1.-** Map of Andalusia (southern Spain) showing the 14 study olive farms (green dots). Dark shaded area in the map shows the occurrence of olive orchards in Andalusia.

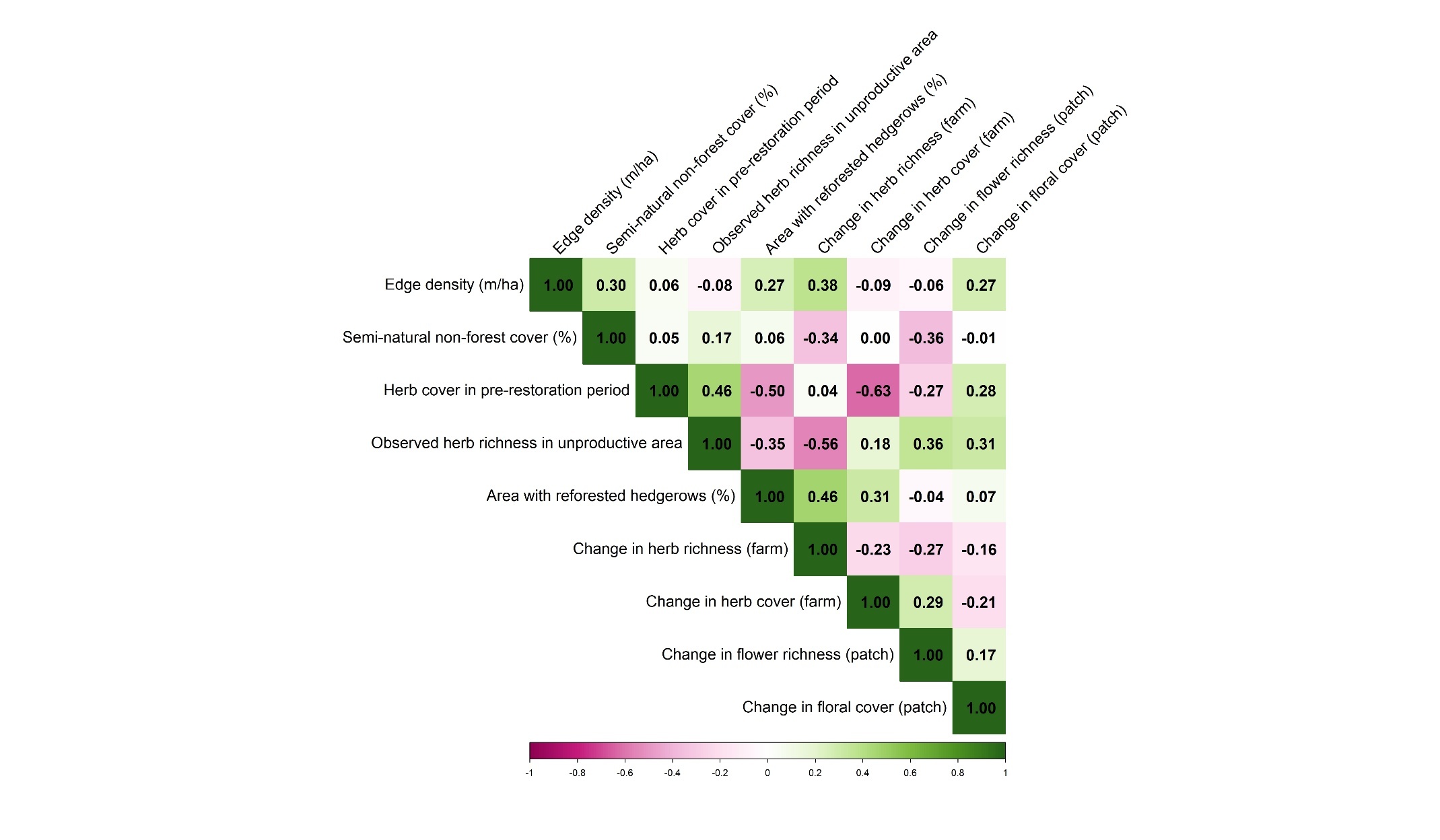

**Figure S2.-** Heatmaps showing the degree of Spearman’s correlation between descriptors of the landscape simplification and the magnitude of change triggered by restoration interventions.
