## Appendix S2 for "Assessing pollinator community recovery in restored agroecosystems using the recovery debt framework"

**Affiliations:**

**Appendix S2: Establishment of the reference system.**

We considered two fundamental criteria in defining the reference systems: landscape heterogeneity and farming intensification. Thus, the overall idea would be that the reference systems should be olive groves (i) within heterogeneous landscapes and (ii) practising low-intensity farming as, based on the literature, they should host the maximum abundance and diversity of bees in olive grove landscapes, and thus be the reference we should approach through restoration.

To look at which of our olive groves could be candidate reference systems, we plotted the study olive groves in two types of scatterplots depicting: (1) the landscape heterogeneity, with two dimensions to account for the compositional (the amount of seminatural habitats) and configurational (edge density) heterogeneity (Martin et al. 2019); (2) farming intensification in olive groves defined also by two dimensions, one considering olive tree plantation density (Morgado et al., 2020; Vasconcelos et al., 2022), separating intensive high-density plantations from traditional low-density tree plantations, and another dimension attending to the percentage of herb cover, which is related to the ground cover management and use of pesticides and that further identified three categories (organic low-intensity ground cover management, non-organic low-intensity ground cover management and high-intensity ground cover management) (Cano et al., 2022; Martínez-Núñez et al., 2019; Martínez-Núñez et al., 2020; Rey et al., 2019). To verify that these dimensions adequately designed reference systems, in these scatterplots we further represented the corresponding value of the bee response variable (abundance and/or richness) in a scale of colour tonality. Candidate reference systems, i.e., those accomplishing better the criteria above, were verified as appropriate reference in each of these scatterplots whenever they reached too the highest (or close to the highest) values of the target variables of bee abundance and species richness.

On the other hand, the wide geographic gradient here considered (mean distance between localities of 80.89 km with a minimum of 9.87 km and maximum of 293.21 km), may configure biogeographic effects on pollinator abundance and diversity that must be accounted for the definition of the reference systems. We conducted three types of analyses to examine how biogeographic noise may influence our target metrics of abundance and diversity. We first carried out a Principal Coordinates Analysis (PCoA) with bee species composition among olive farms and performed a Pearson´s correlation test between the coordinates of PCoA axes for each farm with the geographic coordinates. We further conducted a Mantel test of bee species composition dissimilarity matrices. Finally, we analysed to what extent the variation in abundance and diversity of bees was affected by geographic location using linear models with the geographic coordinates as predictors.

The inspection of the spatial correlation of the variables used to define the reference systems (i.e., richness and abundance of bees) showed that the values of these variables were independent of the geographic location of the olive farms. Spearman´s correlation between PCoA coordinate axes and the geographic coordinates of the olive farms showed that the composition of bee communities did not change across the geographic gradient considered (r < 0.55) (Table S1). A Mantel test also revealed that the dissimilarity in community composition among olive farms did not correlate with geographic distance between localities (r = -0.037; p-value = 0.522). Similarly, linear models showed that the geographic location of olive farms did not affect abundance or species richness values (null models were equivalent to the candidate competing models; see Table S2). These results indicated no need to define separate reference systems according to some geographic stratification. Therefore, any olive farm in our study that satisfies the criteria to be a reference system should be considered.

Scatterplots showed that among the study olive farms under low-intensity ground cover management, only three could be considered candidate reference systems according to their position in the two-dimension heterogeneity and farming intensity spaces (i.e., they were surrounded by a large proportion of semi-natural area, had a high density of edges, high proportion of ground cover and a low density of olive trees). Among these three candidates, two confirmed moderate to high levels of both, abundance and species richness of bees. Hence, we finally chose these two olive farms (marked with the abbreviations “Cas” and “Ta” in the scatterplots of Fig. S1 and S2) as the reference systems for the estimation of RD here.

For convenience, after the validation of candidate reference sites, we designated as the reference state (or value) of the target variable the highest value of the three-year recorded for the chosen candidate farm of pollinators and their function in the reference sites, rather than that of its baseline value, since these reference sites have also been subjected to biodiversity restoration actions. In fact, in the case of diversity indicators at these reference sites, we used the projection to the asymptote of rarefaction curves of species richness as the reference value.

**Table S1.-** Spearman´s correlation coefficient between PCoA axis 1 and 2 (for bee assemblages in floral patches) and geographic coordinates of study olive farms.

|  | UTM_X | UTM_Y |
| --- | --- | --- |
| PCoA axis 1 | 0.01 | 0.06 |
| PCoA axis 2 | 0.55 | 0.42 |

**Table S2.-** Linear models to test the effects of geographic position (geographic coordinates) on the abundance and richness of bees of study olive farms.

| Model code | Fixed factors | AIC | R^2^ |
| --- | --- | --- | --- |
| a) Abundance of bees |  |  |  |
| m_Ab_null | None | 85.941 | 0.000 |
| m_Ab_1 | UTM_X | 87.591 | -0.056 |
| m_Ab_2 | UTM_Y | 87.813 | -0.073 |
| m_Ab_3 | UTM_X + UTM_Y | 89.591 | -0.152 |
| b) Estimated bee species richness |  |  |  |
| m_Ri_null | None | 97.923 | 0.000 |
| m_Ri_1 | UTM_X | 99.370 | -0.041 |
| m_Ri_2 | UTM_Y | 99.034 | -0.016 |
| m_Ri_3 | UTM_X + UTM_Y | 100.990 | -0.106 |

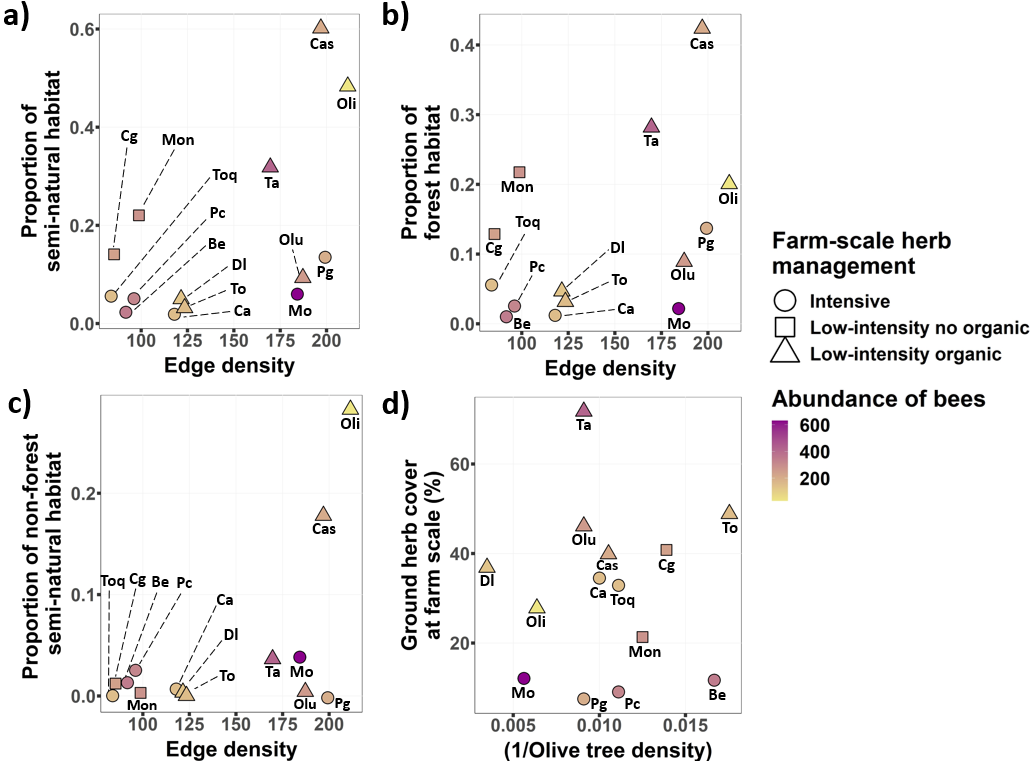

**Figure S1.-** Scatterplot showing the position of each study olive farm in terms of their degree of agricultural intensification considering different variables (i.e., the proportion of semi-natural habitat, the proportion of forest habitat, the proportion of non-forest habitat and edge density, the ground herb cover at farm scale and the olive tree density). Each dot depicts an olive farm filled with a colour scale based on the total abundance of bees on each olive farm. The dots' shape indicates the olive farms' ground herb cover.

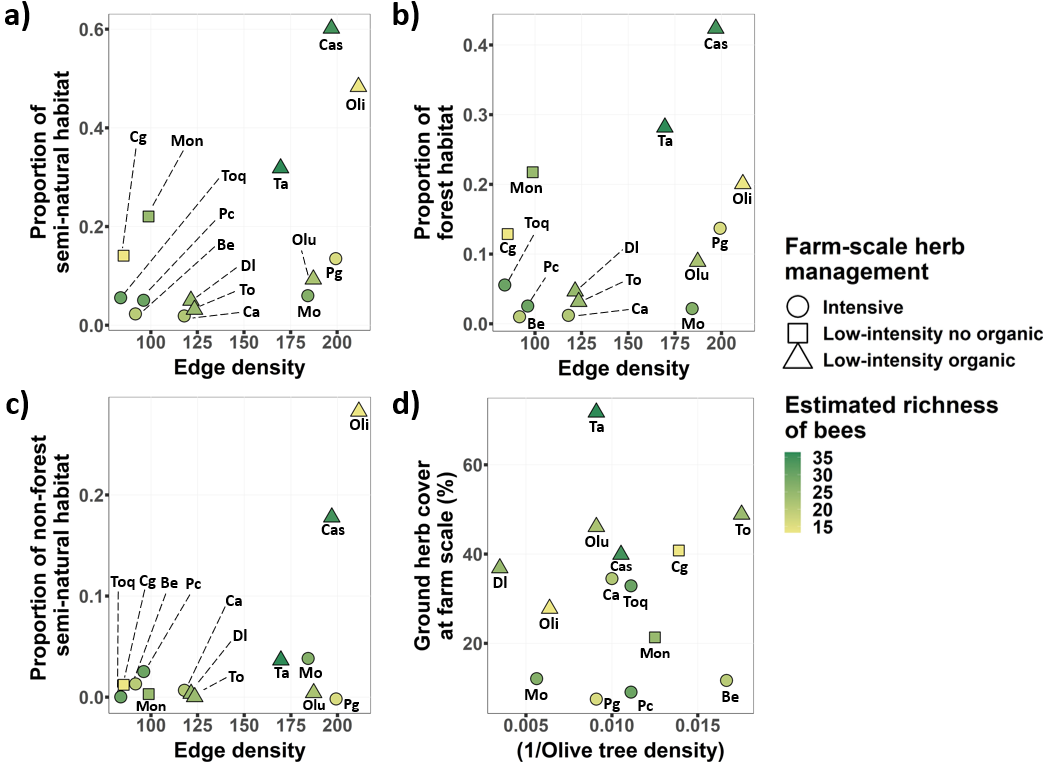

**Figure S2.-** Scatterplot showing the position of each study olive farm in terms of their degree of agricultural intensification considering different variables (i.e., the proportion of semi-natural habitat, the proportion of forest habitat, the proportion of non-forest habitat and edge density, the ground herb cover at farm scale and the olive tree density). Each dot depicts an olive farm filled with a colour scale based on the estimated bee richness of bees on each olive farm. The dots' shape indicates the olive farms' ground herb cover.
