## Appendix S3 for "Assessing pollinator community recovery in restored agroecosystems using the recovery debt framework"

**Affiliations:**

**Appendix S3: Sensitivity analysis excluding and including the reference system from the main statistical tests.**

In this appendix, we compare the main outputs of this study when (a) excluding and (b) including the selected reference farms also in the test sample. The tables and figures show largely consistent results between both situations. Therefore, including the reference sites as test samples, which over time could also approach the reference value (the maximum value for an ecosystem attribute of interest), does not alter the conclusions.

**Section 1) Comparison of the RD between periods (tests in Fig. 2 of the main text) to show the reduction of the RD:**

a) Results excluding the chosen reference farm from the test sample in the analysis:

| **Target variable** | **Factor** | **Mean** | ***P*-value** |
| --- | --- | --- | --- |
| Abundance of bees in floral patches | 2018-2020 | 71.025 | 0.048 |
|  | 2020-2022 | 59.247 |  |
| Bee species richness | 2018-2020 | 51.66 | 0.150 |
|  | 2020-2022 | 40.124 |  |

b) Original table, incorporating the chosen reference farm as a test sample:

| **Target variable** | **Factor** | **Mean** | ***P*-value** |
| --- | --- | --- | --- |
| Abundance of bees in floral patches | 2018-2020 | 71.025 | 0.025 |
|  | 2020-2022 | 55.048 |  |
| Bee species richness | 2018-2020 | 51.66 | 0.100 |
|  | 2020-2022 | 38.573 |  |

**Section 2) Comparison of per year RD for the sets of abundance, diversity and interaction network between categories of original (pre-intervention) herb cover management and use of pesticides in olive farms:**

a) Results excluding the chosen reference farm from the test sample in the analysis:

| **Target variable** | **Category of agricultural intensification** | **Factor** | **Mean ± SE** | **Median** | ***U*** | ***P*-value** |
| --- | --- | --- | --- | --- | --- | --- |
| Abundance of bees in floral patches | Herb cover management | High-intensity | 58.303 ± 7.90 | 62.207 | 14 | 0.370 |
|  |  | Low-intensity | 69.035 ± 5.88 | 68.228 |  |  |
|  | Use of pesticides | Non-organic | 57.708 ± 6.08 | 62.207 | 8 | 0.093 |
|  |  | Organic | 74.281 ± 6.25 | 69.229 |  |  |
| Bee assemblages´ species richness | Herb cover management | High-intensity | 43.623 ± 6.18 | 42.187 | 20 | 0.950 |
|  |  | Low-intensity | 44.775 ± 5.49 | 43.780 |  |  |
|  | Use of pesticides | Non-organic | 46.024 ± 5.11 | 43.006 | 26 | 0.440 |
|  |  | Organic | 41.395 ± 6.67 | 35.177 |  |  |
| Flower visitation rate | Herb cover management | High-intensity | 54.415 ± 8.87 | 63.731 | 13 | 0.792 |
|  |  | Low-intensity | 63.274 ± 8.78 | 61.205 |  |  |
|  | Use of pesticides | Non-organic | 54.277 ± 6.33 | 61.250 | 8 | 0.315 |
|  |  | Organic | 67.944 ± 5.28 | 64.694 |  |  |
| Connectance | Herb cover management | High-intensity | 44.499 ± 12.18 | 26.812 | 9 | 0.329 |
|  |  | Low-intensity | 58.971 ± 7.97 | 62.326 |  |  |
|  | Use of pesticides | Non-organic | 49.592 ± 9.34 | 51.304 | 10 | 0.527 |
|  |  | Organic | 57.294 ± 11.66 | 56.352 |  |  |
| Diversity of interactions | Herb cover management | High-intensity | 18.197 ± 1.62 | 18.050 | 14 | 0.930 |
|  |  | Low-intensity | 20.977 ± 3.45 | 21.043 |  |  |
|  | Use of pesticides | Non-organic | 19.248 ± 2.18 | 18.050 | 14 | 1.000 |
|  |  | Organic | 20.529 ± 4.34 | 21.043 |  |  |

b) Original table, incorporating the chosen reference farm as a test sample:

| **Target variable** | **Category of agricultural intensification** | **Factor** | **Mean ± SE** | **Median** | ***U*** | ***P*-value** |
| --- | --- | --- | --- | --- | --- | --- |
| Abundance of bees in floral patches | Herb cover management | High-intensity | 58.303 ± 7.90 | 62.207 | 20 | 0.662 |
|  |  | Low-intensity | 61.514 ± 9.08 | 66.956 |  |  |
|  | Use of pesticides | Non-organic | 57.708 ± 6.08 | 62.207 | 16 | 0.344 |
|  |  | Organic | 63.378 ± 12.04 | 68.729 |  |  |
| Bee assemblages´ species richness | Herb cover management | High-intensity | 43.623 ± 6.18 | 42.187 | 25 | 0.949 |
|  |  | Low-intensity | 41.867 ± 5.57 | 39.478 |  |  |
|  | Use of pesticides | Non-organic | 46.024 ± 5.11 | 43.006 | 33 | 0.282 |
|  |  | Organic | 38.082 ± 6.37 | 34.150 |  |  |
| Flower visitation rate | Herb cover management | High-intensity | 54.415 ± 8.87 | 63.731 | 18 | 1.000 |
|  |  | Low-intensity | 55.501 ± 8.78 | 61.160 |  |  |
|  | Use of pesticides | Non-organic | 54.277 ± 6.33 | 61.250 | 15 | 0.755 |
|  |  | Organic | 56.128 ± 12.50 | 61.160 |  |  |
| Connectance | Herb cover management | High-intensity | 44.499 ± 12.18 | 26.812 | 11 | 0.343 |
|  |  | Low-intensity | 55.109 ± 7.77 | 51.304 |  |  |
|  | Use of pesticides | Non-organic | 49.592 ± 9.34 | 51.304 | 14 | 0.638 |
|  |  | Organic | 52.222 ± 10.36 | 38.034 |  |  |
| Diversity of interactions | Herb cover management | High-intensity | 18.197 ± 1.62 | 18.050 | 19 | 0.876 |
|  |  | Low-intensity | 18.556 ± 3.79 | 15.498 |  |  |
|  | Use of pesticides | Non-organic | 19.248 ± 2.18 | 18.050 | 21 | 0.638 |
|  |  | Organic | 17.228 ± 4.71 | 15.498 |  |  |

**Section 3) Spearman correlations between the RD of each target variable and the three groups of intensification and the level of change created by restoration:**

a) Results excluding the chosen reference farm from the test sample in the analysis:

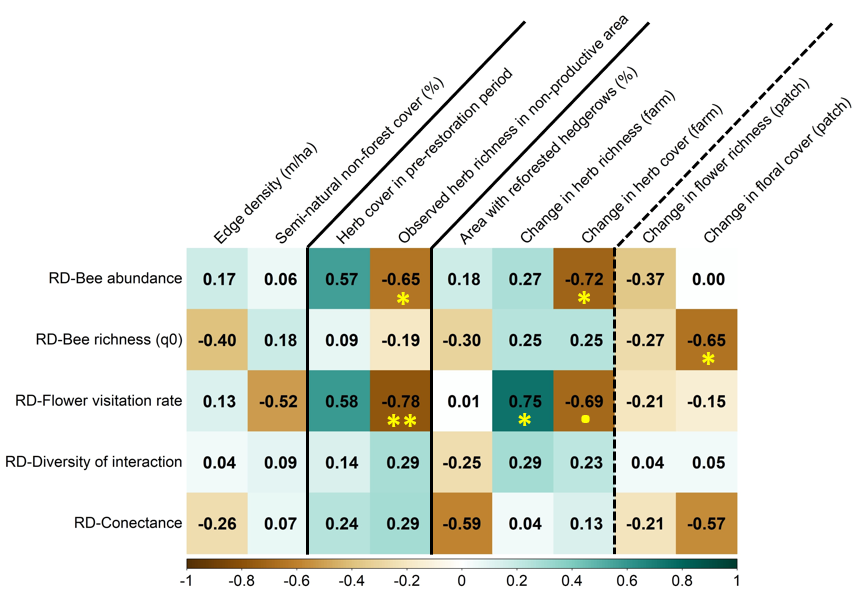

b) Original Figure, incorporating the chosen reference farm as a test sample:

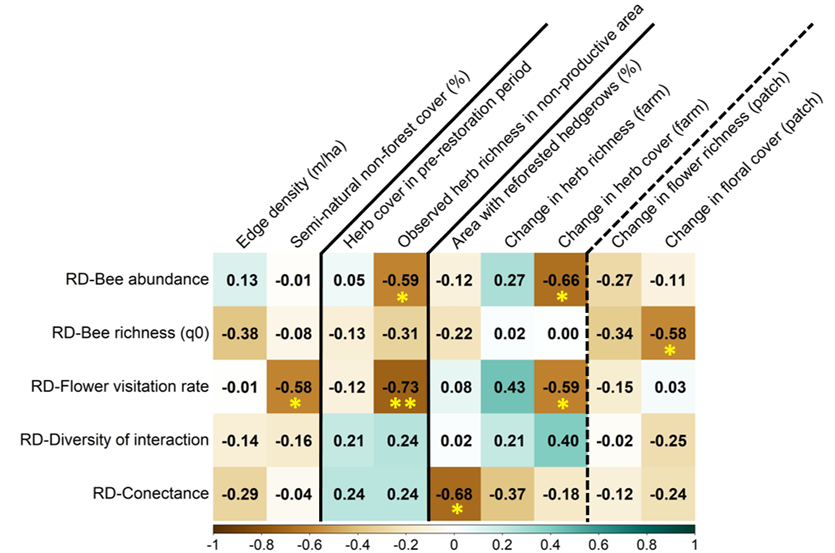

Note that excluding the reference system from the test sample did not change the results qualitatively, either in terms of overall shortening of the recovery debt (section 1 and 2) or in the exploration of the correlates (descriptors of landscape simplification, original ground management (i.e., intensification of the farms), and magnitude of changes triggered by restoration interventions: section 3) of the RD still remaining at the end of the study.
